# A novel model for the induction of postnatal murine hip deformity

**DOI:** 10.1101/270181

**Authors:** Megan L. Killian, Penny R. Atkins, Ryan C. Locke, Michael G. James, Andrew E. Anderson, John C. Clohisy

## Abstract

Acetabular dysplasia is a recognized cause of hip osteoarthritis (OA). A paucity of animal models exists to investigate structural and functional changes that mediate morphology of the dysplastic hip and drive the subsequent arthritic cascade. Utilizing a novel murine model, this study investigated the role of surgically-induced unilateral instability of the postnatal hip on the initiation and progression of acetabular dysplasia and impingement up to 8-weeks post-injury. Specifically, C57BL6 mice were used to develop titrated levels of hip instability (mild, moderate, severe, and femoral head removal) at 3-weeks of age, a critical time for hip maturation. Joint shape, acetabular coverage, histomorphology, immunohistochemistry, and statistical shape modeling were used to assess overall quality of joint health and three-dimensional hip shape following 8 weeks of titrated destabilization. This titrated approach included mild, moderate, severe, and complete instability via surgical destabilization of the murine hip. Acetabular coverage was reduced following severe, but not moderate, instability. Moderate instability induced lateralization of the femoral head without dislocation, whereas severe instability led to complete dislocation and formation of pseudoacetabula. Mild instability did not result in statistically significant morphological changes to the hip. Complete destabilization via femoral head removal led to reduced joint space volume and reduced bone volume ratio in the remnant proximal femur. Collectively, these results support the notion that hip instability, driven by loss of function, leads to morphometric changes in the maturing mouse hip. This model could be useful for future studies investigating the mechanical and cellular adaptations to hip instability during maturation.

## Introduction

The rising rate of osteoarthritis (OA) in young adult patients (i.e., younger than 50 years of age) is a growing concern.^1–3^ It is now widely believed that structural hip disease, including acetabular dysplasia and femoroacetabular impingement (FAI), are the predominant etiologies of hip OA, especially among younger adults.^4–7^ Acetabular dysplasia, which is characterized by insufficient femoral head coverage, has been implicated as the cause of hip OA in 20-40% cases; however, FAI, which is characterized by an excessively-prominent acetabular rim and/or an abnormally-shaped proximal femur, may account for up to 82% of hip OA cases. ^7,8^ Damage to the cartilage and labrum in patients with acetabular dysplasia and FAI suggests that these conditions initiate OA through altered chondrolabral (i.e. cartilage and labral) mechanics as well as abnormal kinematics. Dysplastic hips present with joint laxity and instability ^9,10^, and studies suggest FAI patients have limitations in their range of motion. Further, computational studies have demonstrated that acetabular dysplasia patients have higher cartilage contact stresses ^11–13^, and increased load transfer to the labrum ^14^. Similar modeling studies have shown increased shear stresses to the cartilage of FAI patients ^15–17^.

It is important to recognize that acetabular dysplasia and FAI are not mutually exclusive.^18,19^ Specifically, the dysplastic hip has been characterized as having an elliptical shaped femoral head,^19,20^ increased femoral neck waist,^19^ and decreased epiphyseal extension towards the femoral neck.^19^ In addition, patients with cam-type FAI can present with loss of femoral coverage ^21,22^, and patients with pincer-type FAI due to acetabular retroversion have deficient posterior coverage ^23^. However, it is still unclear what drives acetabular or femoral malformations during adolescent growth, and how these pathological adaptations influence cartilage and bone health of the hip.

To better understand the mechanisms by which pathological adaptations occur in the hip, we and others have explored the normal development of joint formation, growth, and maturation.^24–29^ It is generally accepted that hip joint formation and maturation depends on a well-orchestrated series of molecular and mechanical events throughout the duration of perinatal development.^28,30^ In mice, maturation of the proximal femur occurs during postnatal development between 7 and 28 days, when a single chondroepiphysis of the proximal femur discretizes into two epiphyses that separate the femoral head and trochanter.^31^ The murine hip reaches maturation by 8 weeks of age, indicated by fusion of the triradiate cartilage of the pelvis that connects the ilium, ischium, and pubis. ^28^ In humans, this maturation occurs during adolescence between 11-18 years of age.^32^ The developmental timeline of the musculoskeletal system, in general, is fairly well characterized for mice and humans. However, there is a need to develop and deploy small animal models to better understand how mechanical loads during musculoskeletal growth are associated with hip pathology.

The purpose of this study was to identify how destabilization of the hip influences joint maturation, shape of the proximal femur, and hip health during skeletal maturation in mice. We hypothesized that destabilization of the murine hip and subsequent changes in loading during postnatal growth, prior to triradiate growth plate fusion, would cause acetabular dysplasia and focal chondrolabral degeneration. To investigate this hypothesis, we developed a novel, titrated model of hip instability in 3-week old neonatal pups. After 8 weeks of instability, we compared bone morphometry, histomorphology, immunohistochemistry, shape adaptations, and bone apposition adaptations in response to each level of hip instability.

## Methods

### Animal Model

All studies were performed in compliance with the Animal Studies Committee at the Department of Comparative Medicine at Washington University. 3-week old C57BL6 mouse pups (N = 36, bred in-house) were used for this study to investigate the role of loading at a key stage of hip maturation. All surgeries were performed by a board certified orthopaedic surgeon (JCC) using a high-resolution dissection microscope under aseptic conditions. Mice were anesthetized (2-3% isofluorane with oxygen carry) and subjected to titrated unilateral surgical destabilization via iliacus/iliopsoas tenotomy, capsulotomy, and/or ligamentum teres transection. Instability was confirmed via acute lateralization of the femoral head with the anesthetized mouse lying in supine position. For each destabilization, a 1-cm incision was made into the ventral midline of the hip. The leg was placed in hip flexion, abduction, and external rotation and held in place by an assistant. Care was taken not to damage the neurovascular bundle. For mild destabilization (N = 4; 3 males, 1 female), a full capsulotomy, iliopsoas tenotomy, and ligamentum teres transection was performed using curved, pointed-tip ophthalmic surgical scissors. Transection of the ligamentum teres was performed after capsulotomy and tenotomy by gentle lateral subluxation for intra-articular exposure of the ligament. For moderate destabilization (N = 10; 5 males and 5 females), capsulotomy, iliopsoas tenotomy, and ligamentum teres transection was performed as it was for mild instability. However, a partial hip abductor tenotomy was also performed by detaching the piriformis, superior and inferior gemellus, and quadratus femoris from its femoral insertion sites. For severe destabilization (N = 12; 7 males and 5 females), the same procedure of moderate instability was performed. However, a full hip abductor tenotomy was also performed by detaching the gluteal rotators (gluteus minimus, obturator internus, gluteus maximus, and gluteus medius) from their proximal femoral insertion sites. Femoral head resection (FHR; N = 7; 5 males and 2 females) was performed to induce complete unloading of the acetabulum for comparison to destabilization groups. For FHR, capsulotomy, iliopsoas tenotomy, and ligamentum teres transection was performed, but all other soft tissue attachments were maintained. The femoral head was cut using ophthalmic surgical scissors and carefully removed from the joint space.

In each destabilization model, the hip was reduced by the surgeon prior to incision closure. The capsule was left unrepaired. Upon completion of soft tissue injuries, the injury site was treated with bupivacaine hydrochloride (Marcaine, Hospira, Lake Forest, IL) and the surgical site was closed using 6-0 Vicryl (Ethicon, Somerville, NJ). A sham operation was performed on the contralateral side that included exposure of the capsule, identification of the hip joint, and closure of the incision without joint (muscle/ligament) damage. Mice were administered a subcutaneous injection of buprenorphine (Buprenex, Richmond, VA) at a dose of 0.05mg/kg. Following surgery, mice were housed with same-sex littermates (less than 5 per cage) in cages with corn bedding with food and water ad libitum, exposed to a 12hr on/12hr off light/dark cycle, and monitored post-operatively for pain and distress on a daily basis.

### Bone and Joint Morphology

While under isoflurane anesthesia, mice from the mild, moderate, and severe instability groups were scanned in supine position with hips in extension using *in-vivo* micro-computed tomography (VivaCT, Scanco, Switzerland; 21μm voxel, 55kVp, 145μA, 100msec integration time) prior to injury and at 2-, 6-, and 8-weeks post-injury. The region of interest selected for imaging included bilateral hip joints. Three-dimensional CT images were exported as DICOM files for image processing. Acetabular coverage of the proximal femur was measured by using previously described methods. ^28^ Briefly, hips were digitally repositioned and reoriented using the 3D MPR tool in OsiriX (64-bit software, Pixmeo, Geneva, Switzerland). Norberg angles (α) were evaluated for both hips in frontal plane ^28^.

At 8-weeks post-injury, mice were sacrificed, eviscerated, and the hip joint complex was isolated intact with surrounding musculature removed for higher contrast imaging using *ex-vivo* microCT (microCT40, Scanco, Switzerland; 20μm voxel size, 70kVp, 114μA, 200msec integration time). Three-dimensional CT images were exported as DICOM files and analyzed using CTan (Bruker, Kontich, Belgium) and FIJI/ImageJ.^33^ Using CTan, regions of interest (ROI; pelvis, proximal femur, and acetabular joint space) were manually traced based on anatomical landmarks, with interpolation between every 10^th^ slice. Landmarks were chosen that were consistent across all specimens (Supplemental Figure S1). The number of slices per ROI varied as size and distance between chosen landmarks differed between samples. The ROI used to determine pelvis bone morphometry was defined as the entire tissue volume of the acetabulum, beginning at 20 consecutive slices proximal to the femoral head (FH), and continued distally until the connection of the ischium to the pubis bone was observed. The ROI for the proximal femur included the entire volume of the FH and neck. The intertrochanteric line was used as the distal boundary of the femoral neck, and the greater and lesser trochanters were excluded from femoral neck measurements. Acetabular joint space was measured from the proximal acetabular rim to the acetabular notch using a contour of the lunate surface, acetabular fossa, and an interpolated plane of the acetabular rim in consecutive CT images. The volume of the acetabular joint space inside this plane was measured using BoneJ’s Moments 3D plugin.^34^ Visualization of morphology of each traced region was performed using 3D reconstructions generated in OsiriX MD 8.0.1 (Pixmeo, Switzerland) and MeshMixer (Autodesk, California, version 3.0). Gaussian blur (kernel: square, radius: 0.5), unsharp mask (kernel: square, radius: 2, amount: 200, threshold: 0), multi-level Otsu thresholding (levels: 5, threshold: level of class), and adaptive thresholding (type: median, kernel: square, radius: 4, threshold: 90-255) was performed to segment bone from background noise prior to quantitative analysis of total volume (TV), bone volume (BV), bone volume fraction (BV/TV), and acetabular joint space volume (JSV). TV and acetabular JSV were measured as all pixels inside the contour (and excluding pixels in contact with contour line).

### Dynamic Histomorphology

Bone formation and morphology was assessed using a subset of mice (N = 5, 3-week old C57BL6, 4 males and 1 female) from the severe instability group. Age-matched littermates (N = 3, all females) were used as uninjured controls. Mice were administered calcein green (10mg/kg, Sigma-Aldrich, St Louis, Missouri) and alizarin complexone (30mg/kg, Sigma-Aldrich, St Louis, Missouri) by intraperitoneal injection at 10 days and 3 days prior to death, respectively. Mice were euthanized 8 weeks after instability surgery. Right and left femur were disarticulated from the joint and fixed in 10% formalin overnight and stored in 70% ethanol (4°C), until microCT scanning (microCT40, 20μm resolution, 70kV, 114mA, 200msec integration). Samples were then dehydrated in serially increasing ethanol concentrations (70-100%) followed by xylene, and then infiltrated and embedded in polymethylmethacrylate. Three to five sections (85-105μm thick) per femur were cut in the frontal plane using a microtome (Leica SP 1600, Buffalo Grove, IL, USA). Sections were mounted on glass slides, ground to 30-40μm thickness, and imaged using a fluorescent microscope (Zeiss Apotome.2, Thornwood, NY, USA) using green fluorescence protein (GFP) and TexasRed filters for calcein and alizarin visualization, respectively. Images were digitized, stitched in Zen2 software (Zeiss, Thornwood, NY, USA), and qualitatively assessed for bilateral bone formation.

### Histology and Immunohistochemistry

Following *ex-vivo* microCT, hips were decalcified in formic acid solution (ImmunoCal, American MasterTech, Lodi, CA, USA) for 48-72 hours. After decalcification, tissues were processed for paraffin embedding and sectioned in frontal plane at 5-micrometer thickness. Sections were stained with safranin o/fast green with Weigert’s hematoxylin for cartilage quality, extracellular matrix morphology, and cellular localization. Stained sections were imaged on Aperio AT2 ScanScope (Leica Biosystems, Wetzlar, Germany), and Aperio ImageScope (Leica Biosystems, Wetzlar, Germany, Version 12.3.0.05056) was used to obtain 5x images of the entire hip joint and 20x images of the articular cartilage of the FH and acetabulum.

Immunolocalization of collagen type II (COL2) and matrix metalloproteinase-13 (MMP13) was also performed on paraffin sections. Three biological replicates from each group (Mild, Moderate, Severe, and FHR) were used. Sections were blocked by 5% goat serum and incubated with primary antibodies (COL2: rabbit polyclonal, Abcam ab34712; MMP13: mouse monoclonal; Santa Cruz sc515284) at 1:100 dilutions. For negative controls, sections were incubated with blocking buffer without primary antibodies. Antibody detection was performed using a commercially available kit (DAB peroxidase substrate kit, Millipore). Sections were counterstained with Mayer’s hematoxylin and imaged using light microscopy (Zeiss, Gena, Germany).

### Statistical Shape Modeling

MicroCT image volumes of bilateral mouse hips from the severe instability group (N = 7; 4 males and 3 females) were segmented and reconstructed in Amira (FEI 5.6 Merignac, France) for statistical shape modeling. The surfaces were scaled isotropically and smoothed to remove segmentation artifacts. Surfaces were reflected, as necessary, and aligned using the iterative closest point algorithm implemented in Amira.

Using preprocessing tools for ShapeWorks, distance transforms were generated for the scaled femur surfaces to define the femur shape within the image volume implicitly. ^35^ An iterative splitting and optimization routine implemented in ShapeWorks was used to automatically place correspondence particles (n=2048), avoiding the need for manual particle placement or the use of training shapes. ^35^ A plane located below the third trochanter for each femur was used to limit the correspondence particles to the proximal femur.

The high-dimensional data of the correspondence particle locations was reduced to a smaller set of linearly uncorrelated components (modes) used to describe the variation of the shapes using principal component analysis (PCA). Mean femur surfaces for the injured and contralateral femurs were generated from the correspondence points and were smoothed and decimated using algorithms in PreView (Version 1.18.1).^36^ Morphological differences between shapes were calculated as the distance between the mean shape of the severe instability and mean shape of the contralateral femurs.

### Statistical Analysis

Statistical comparisons of Norberg angle and bone morphometry between injured and contralateral hips were compared using a 2-way ANOVA (groups: injury, limb) using Prism 6.0h (Graphpad, LaJolla, CA). Specifically, differences in Norberg angle between contralateral and injured hips (side) for the severe instability group at 2-, 6-, and 8-wks post-injury were compared using a 2-way ANOVA, matching for time and side, using a Sidak’s multiple comparisons test. Comparisons between instability models at 8wks post-injury were performed for the following outcomes: JSV, proximal femur BV, TV, and BV/TV, and pelvis BV, TV, and BV/TV. Data were compared using 2-way ANOVA with repeated measures (side) with Sidak’s multiple comparisons tests.

For statistical shape modeling, the significant (non-spurious) modes of variation were identified using parallel analysis of the principal component loading values. Overall shape difference between the severe instability and contralateral femurs were identified using Hotelling’s T-squared test. The specific modes of variation which represented significant shape differences were determined using a Student’s T-test. Statistical analysis of shape variation was completed in R.^37^

## Results

### Acetabular Coverage Patterns and Hip Morphometry

Most pups tolerated the instability surgical procedures well, with one fatality at time of operation (mild instability group, female). Mild instability did not lead to noticeable changes in FH coverage (Figure 1A), Norberg angle (Figure 3), or qualitative bone shape at 2-, 6-, or 8-weeks post-injury. For mice subjected to moderate instability, lateralization of the proximal femur (Figure 1B) with decreased Norberg angle (Figure 3) was observed. For mice subjected to either severe instability or FHR, Norberg angle and FH coverage were not calculated due to complete loss of FH in the acetabulum for severe (Figure 1C) and FHR (Figure 1D) groups.

**Figure 1.**
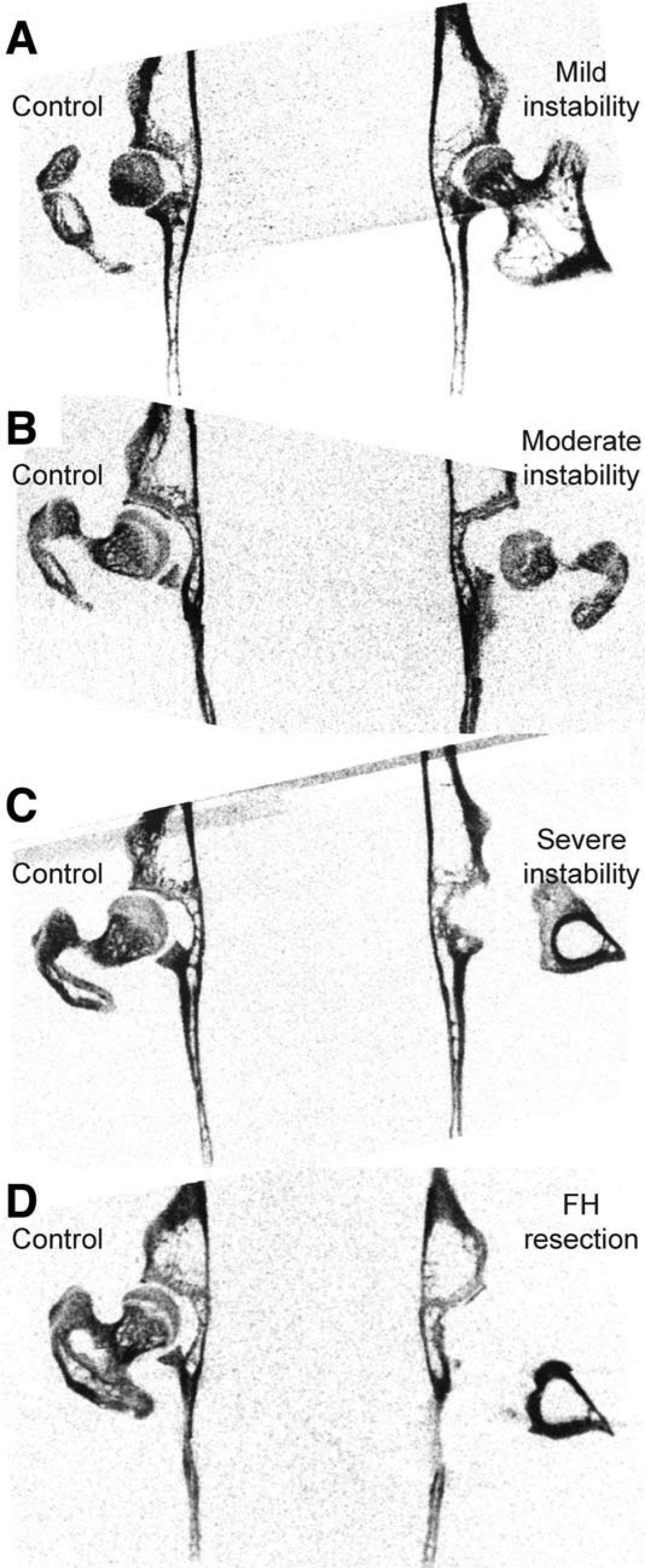
Two-dimensional (2D) planar x-ray images (1mm cut plane thickness) from reconstructed microCT of representative bilateral hips for (A) mild instability, (B) moderate instability, (C) severe instability, and (D) FH resection groups at 8-weeks post-surgery. Contralateral (control), sham-operated hips (left) appeared morphologically similar between groups.

Moderate destabilization did not lead to noticeable three-dimensional morphological changes to the proximal femur or pelvis at 8-weeks post-injury (Figure 2A). After 8-weeks of severe instability, however, the proximal femur in the injured hip appeared to have more radio-opaque bone compared to the sham-operated, contralateral side (Figure 2B). Additionally, severe instability resulted in closure of the acetabulum and a rounded acetabular rim (Figure 2C). Proximal femurs of the FHR group were non-existent following surgical removal and the remaining femoral neck was rounded and mineralized (Figure 2C). The pelvis demonstrated a similar morphology of acetabular closure and rounded acetabular rim compared to the severe instability group (Figure 2C).

**Figure 2.**
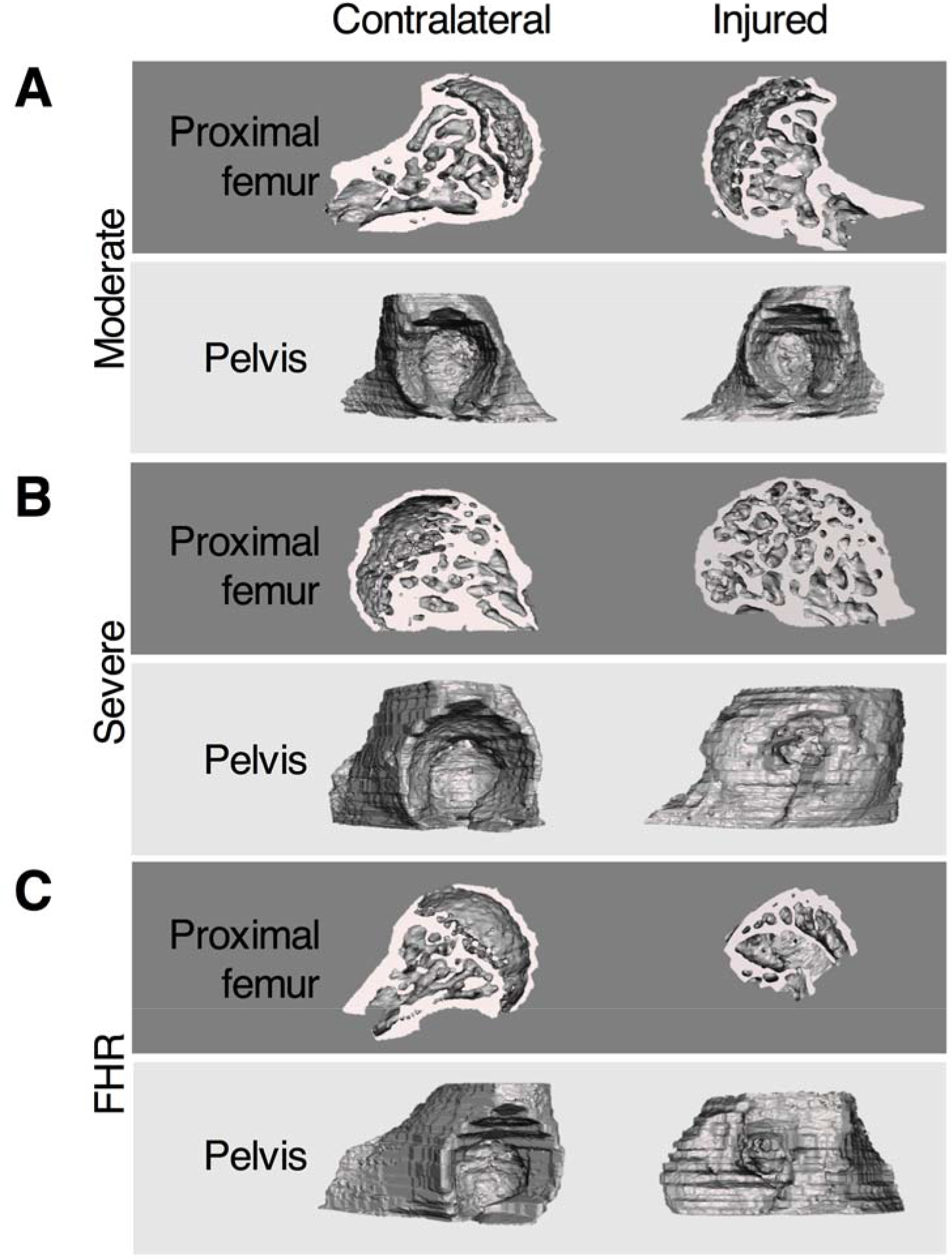
MicroCT reconstructions of the proximal femur (frontal cut plane) and pelvis (sagittal view plane) for contralateral and injured hips in the (A) moderate destabilization, (B) severe destabilization, and (C) FHR groups.

**Figure 3.**
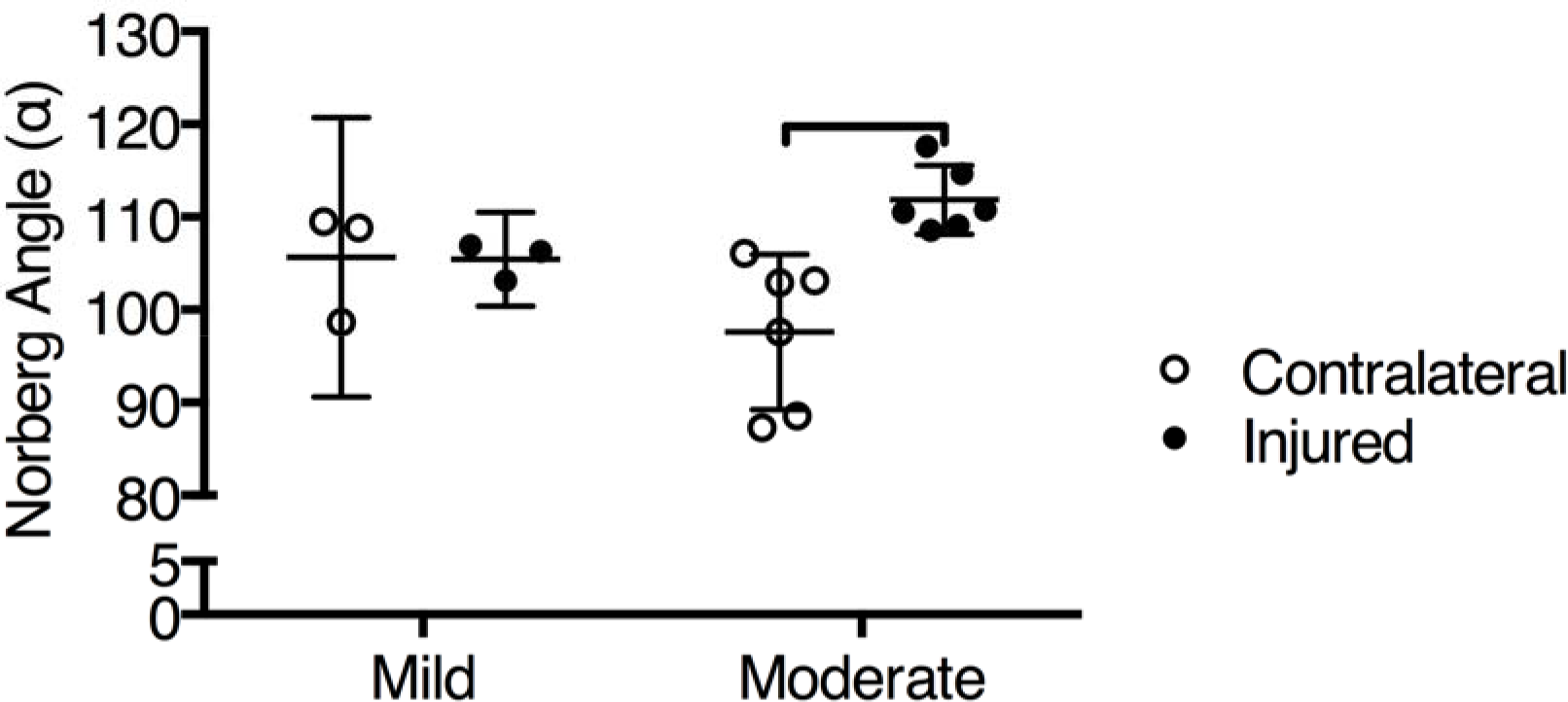
Norberg angle measurements for mild and moderate destabilization (injured) hips compared to contralateral hips measured using vivaCT and Osirix, at 8-weeks post-injury. Bar indicates statistically significant difference (p < 0.05) between contralateral and injured hips within injury group. Data presented as mean ± 95% confidence interval.

Pelvis TV was significantly increased following FHR compared to contralateral hips (Figure 4A). However, no significant differences in BV were observed between injured and contralateral pelvi for either moderate or severe injury groups (Figure 4B). FHR led to significantly increased pelvis BV/TV of the contralateral hip compared to the injured hip (Figure 4C), as well as compared to contralateral pelvi of the moderate and severe instability groups (Figure 4C). FHR also resulted in reduced proximal femur TV of the injured hips compared to contralateral hips (Figure 4D). However, no differences in femur TV were observed for moderate or severe hips compared to their paired contralateral sides (Figure 4D). For the FHR group, BV was significantly decreased in proximal femur compared to contralateral hips (Figure 4E), and a significant decrease in proximal femur BV was observed in FHR compared to the same region in severe instability (Figure 4E). No significant differences were observed in proximal femur BV/TV between or within any groups at 8-weeks post-injury (Figure 4F).

**Figure 4.**
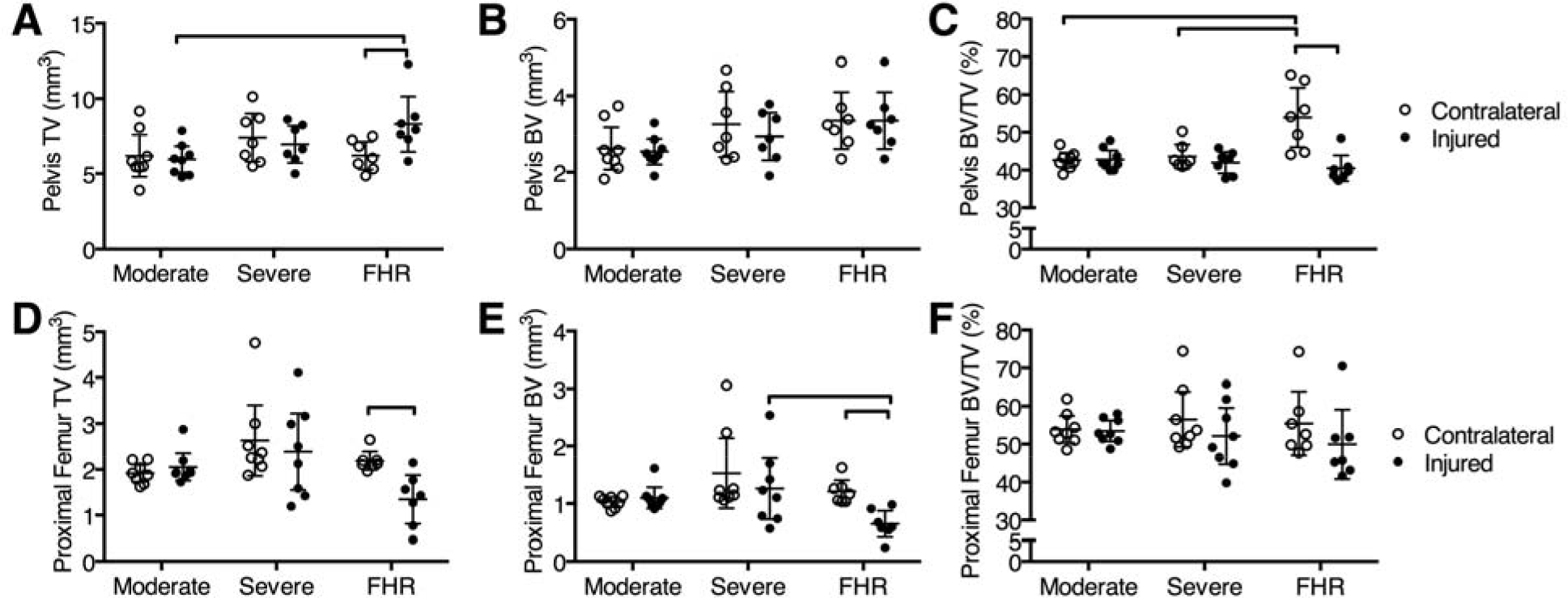
Bone morphometric outcomes of the (A-C) pelvis and (D-F) proximal femur of moderate, severe, and FHR contralateral and injured hips, including (A & D) total volume (TV), (B & E) bone volume (BV), and (C & F) bone volume/total volume ratio (BV/TV, %). Lines indicate significant difference between respective limbs or within group (contralateral vs. injured), p < 0.05. Data presented as mean ± 95% confidence interval.

Acetabular JSV was significantly reduced at 8-weeks post-injury for both severe instability and FHR compared to contralateral hips (Supplemental Figure S2). Additionally, JSV was significantly reduced for both severe instability and FHR compared to moderate instability (Supplemental Figure S2). JSV showed no significant change following injury in the moderate instability group compared to contralateral hips (Supplemental Figure S2).

### Microscopic Hip Joint Changes

No distinguishable differences were observed histologically between contralateral and injured hips following mild instability, and cartilage appeared healthy and intact on both the femoral head and acetabulum (Figure 5A-B). Moderate instability resulted in bilateral asymmetry in the formation of secondary ossification center, with injured hips demonstrated increased bone formation compared to contralateral hips (Figure 5C-D). Articular cartilage of the femoral head and acetabulum appeared intact and healthy in both contralateral and moderate instability hips (Figure 5C-D). Severe instability led to abnormal femoral head and trochanter shape, as well as formation of a pseudoacetabulum (Figure 5F). Following severe instability, fibrocartilage growth was observed on the proximal diaphysis/lesser trochanter of the femur, at points where the pelvis and femur articulate (Figure 5F & F’). Additionally, cells at the articulating surface between femur and pseudoacetabulum had a rounded, fibrochondrocyte-like morphology (Figure 5F’). FHR led to formation of cartilage-like tissue on the residual femoral neck (Figure 5H & H’). Additionally, the thickness of cartilage present appeared thickened for both residual femur and pseudoacetabulum on injured FHR hips (Figure 5H’). Heterogeneous distribution of red stain, indicating sulfated glycosaminoglycans, were also present at the femur/pseudoacetebular interface (Figure 5H & H’). Contralateral hips for moderate, severe, and FHR hips appeared normal with intact cartilage, rich proteoglycan stain, and intact articular surfaces (Figure 5A, C, E, and G). Following 8-weeks of severe destabilization, the femoral head appeared less mature and less mineralized compared to the contralateral femur, as shown using dynamic histomorphometry (Supplemental Figure S3).

**Figure 5.**
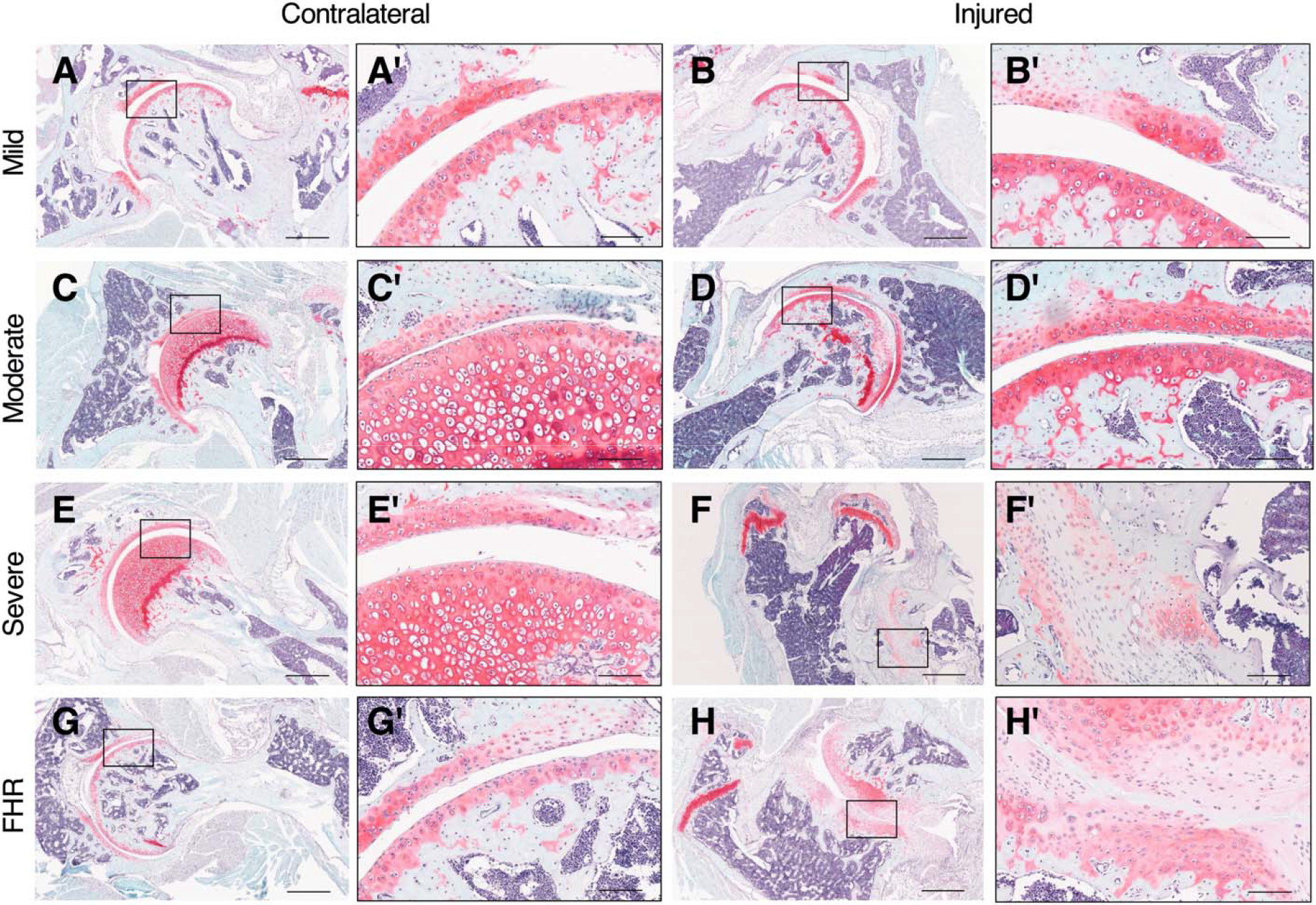
Histological sections stained with Safranin O/Fast Green FCF/Weigert’s Hematoxylin for contralateral and injured hips of Mild (A & B), Moderate (C & D), and Severe (E &F) instability as well as FHR (G & H) groups. High magnification (A’-H’) images highlight the inset boxed region in the respective low magnification images (A-H). Scale bar = 500μm for A-H; 100μm for A’-H’.

The articular cartilage of the femoral head showed similar localization of COL2 between contralateral and injured hips for the Mild and Moderate groups (Figure 6A&B; Figure 6F&G). COL2 was distributed throughout the pseudo-acetabulum of the injured hip from both Severe and FHR (Figure 6P) groups. COL2 was absent in the articular cartilage of the injured hip from the Severe group (Figure 6L). MMP13 was localized in the articular cartilage of contralateral, but not injured, hips in the Mild, Moderate, and Severe groups (Figure 6C-N). MMP13 was localized in the pseudo-acetabulum of both the Severe and FHR groups (Figure 6R) and was not prevalent in articular cartilage of FHR contralateral hips (Figure 6Q). Positive staining for COL2 and MMP13 was not observed in negative control sections Figure 6E&J, respectively).

**Figure 6.**
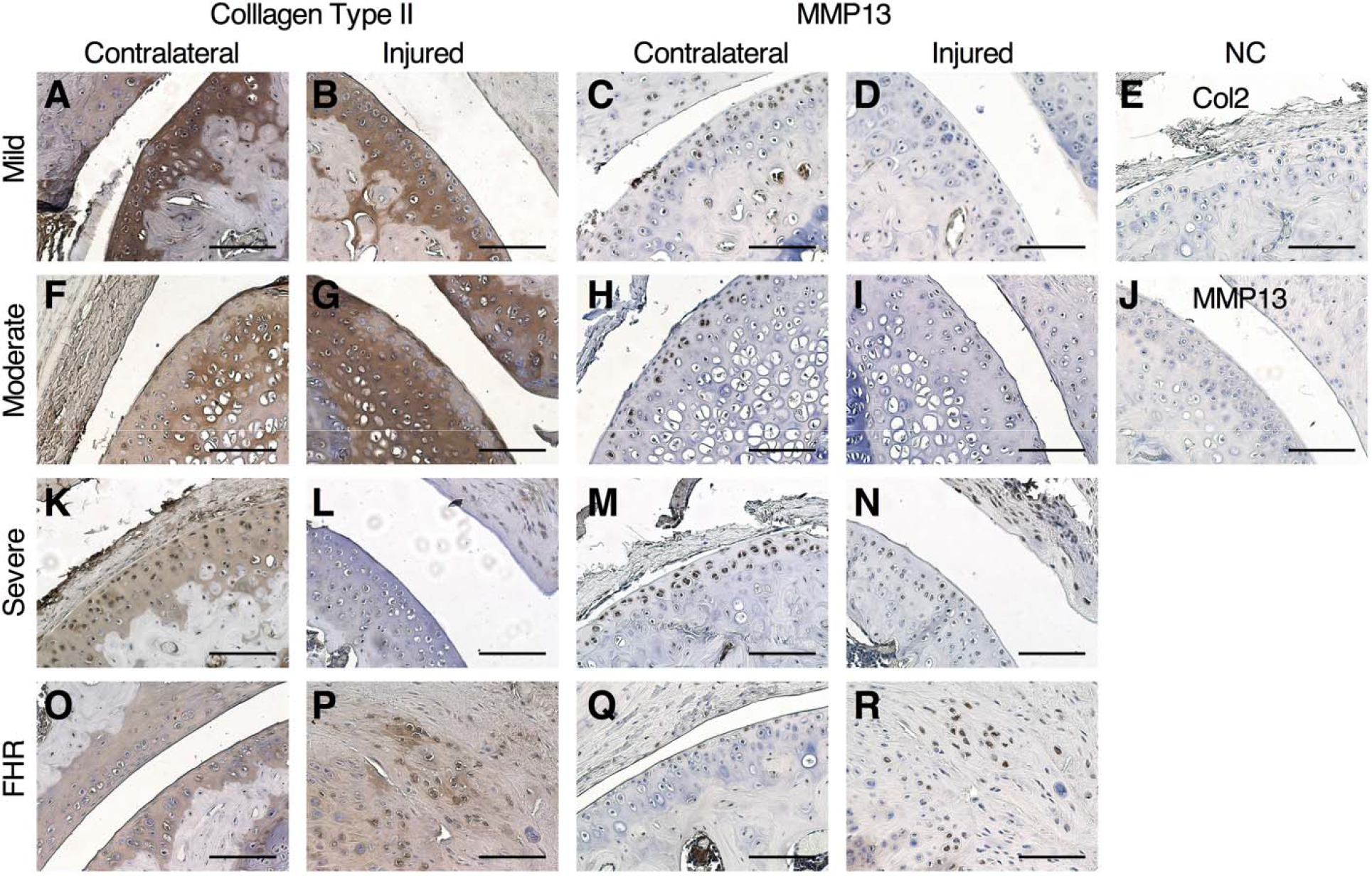
Immunohistochemistry for COL2 and MMP13 following titrated instability. COL2 localization of the (A-O) Contralateral and (B-P) Injured limb for the (A&B) Mild, (F&G) Moderate, (K&L) Severe, and (O&P) FHR groups. MMP13 localization of the (C-Q) Contralateral and (D-R) Injured limb for the (C&D) Mild, (H&I) Moderate, (M&N) Severe, and (Q&R) FHR groups. The negative control slides for (E) COL2 and (J) MMP13 immunohistochemistry. Scale bar for all images: 100μm. Images taken at 20x magnification.

### Changes in Proximal Femur Shape

For statistical shape modeling, the first five modes of variation from PCA accounted for 90% of the population variation; however only the first three were determined to be significant, accounting for 80.1% of the shape variation. Of these three modes, the first contained 47.2%, the second 24.2%, and the third 8.8% of the total variation. Only mode three represented significant shape differences between the injured and contralateral hip (p=0.02), specifically describing the variation in neck shaft angle and sphericity of the femoral head, as well as the prominence of the lesser trochanter. The regions of greatest shape difference corresponded to the variations represented by PCA mode three (Figure 7).

**Figure 7.**
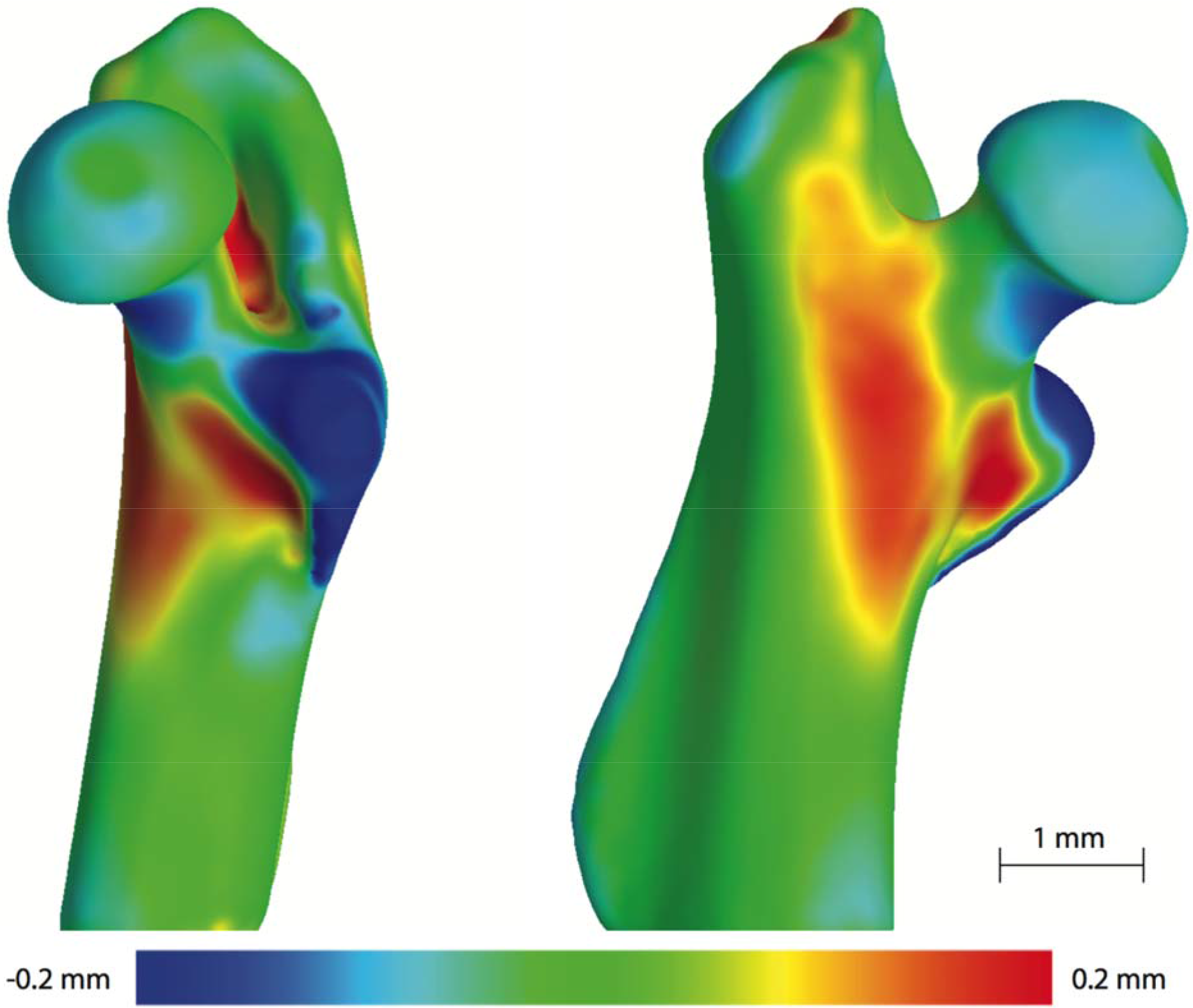
Surface difference between the population-based mean contralateral and severe instability femurs mapped onto the mean contralateral femur. Regions of reduced bone are indicated in blue and increased regions in red. Scale bar = 1 mm.

## Discussion

This is the first study of its kind to identify the role of various magnitudes of hip instability on the postnatal, growing mouse hip. This mouse model is a potential small animal analogue of post-natal hip dysplasia. We developed a model for the controlled destabilization of the hip that resulted in lateralization of the femoral head without dislocation (i.e., moderate destabilization) or severe instability and dislocation (i.e., severe destabilization), with controllable and measurable morphological changes to the proximal femur and acetabulum. Our animal model may aid in identifying more reliable biomarkers or radiographic abnormalities of the hip that can be used in newborn screening to preserve a critical window for intervention. There is ample room for improvement in the clinical management and potential circumvention of hip dysplasia. Acetabular dysplasia is a perplexing problem for clinicians both in terms of managing the unstable hip itself, as well as managing the resultant OA. An animal model that helps identify the biologic and mechanical underpinnings of hip destabilization represents a potential opportunity for the identification of novel diagnostic and treatment modalities.

Murine models have been used for over two decades to study the pathophysiology of knee OA. ^38^ The use of surgical and non-surgical models for instability-driven approaches have pushed the field forward by allowing for the identification of initiation and progression of cartilage damage specific to the knee. Historically, the use of knee injury induced via surgical (destabilization of the medial meniscus ^38^; ligament transection ^39^) or nonsurgical ^40^ trauma has elucidated the importance of joint stability on cartilage health and maintenance. However, a large gap exists for the study of OA development and progression in load-bearing joints other than the knee. Additionally, our understanding of cartilage health and maintenance may be dependent on the anatomical environment and joint-specific considerations for joints other than the knee. To date, there is a dearth of animal models that explore hip cartilage health^28,41,42^, and none that utilize surgical approaches for the induction of postnatal injuries. In this study, we have developed a new surgical injury model for improving understanding of the health and structure of the growing, postnatal hip. We show that soft tissue injury, including ligament transection and tenotomy, resulted in increased cartilage damage, fibrosis, and overall degenerative changes to the joint. The shape of the hip joint was severely affected by the detachment of soft tissue structures during postnatal growth, highlighting the critical role of soft tissue attachments and proper joint loading during the maturation of the hip.

Potential targets include MMP13 and COL2, markers of cartilage remodeling that were explored in this study. The presence of MMP13 is commonly accepted as an indicator of osteoarthritis (OA) ^43,44^. Yet in our titrated model of developmental hip dysplasia, MMP13 was primarily localized to the contralateral hip; this suggests that injury may induce abnormal biomechanics in the non-injured hip, similar to the human scenario of hip dysplasia ^45,46^. In contrast, MMP13 localization to the contralateral hip may indicate that there is homeostasis of COL2 breakdown and synthesis, as no differences in COL2 staining were observed between mild, moderate, and severe groups. It is worth noting that our analysis of MMP13 is a snapshot of time after injury and its localization may differ at an earlier or later time after injury. The severe injured group lost COL2 staining of the articular cartilage, suggesting late stage OA and loss of structural integrity^43^.

This is the first study to quantitatively model the shape of the proximal femur along with quantitative comparisons between normal development and development with morphological deformity in the murine hip. Using statistical shape modeling approaches, we found that the regions of largest shape variation following severe destabilization were localized to the lesser trochanter and intertrochanteric line, and these regions do not directly align with muscle insertion sites that would have been affected by the severe instability injury. Accordingly, alterations in loading due to instability within the joint likely have a more complex effect on bone morphology during development. Previous work by Chan et al. has shown that shape variations exist during normal growth in human patients with morphological deformities of the hip, such as Legg-Calvé-Perthes disease and slipped capital femoral epiphysis. ^47^ Work by this group also demonstrated that the growth and shape attainment of the distal femur are comparable between humans and mice. ^48^ Our present work provides a platform for understanding the progression and development of proximal hip deformities in a small animal model that mimic morphological deformities observed in the clinic, such as LCP, SCFE, and developmental dysplasia.

With the development of this model, we will now be able to explore the role of mechanical loading via surgical hip instability on hip OA and morphological adaptations during post-natal growth and mineralization. For example, future work will be able to pair this surgical model with genetically modified mouse models for lineage tracing and genetic deletion of key molecular targets, such as *gli1* and *sox9*, in order to better understand the mechanobiological regulation, cell proliferation and differentiation, and joint pathology related to perturbed hip maturation during adolescent growth.

### Limitations

Although this model was developed for the induction of hip injury, we did not quantify the magnitude of instability that was induced following soft tissue injury. The effect of this soft tissue injury on long-term stability of the joint may prove relevant for ascertaining the healing and mechanical loading of the hip joint *in-vivo*. For well-established small animal models of knee OA, the quantitative assessment of anterior-posterior and varus-valgus translation are commonly used as methods for assessing stability and laxity of the knee joint. ^49,50^ However, at this time, no such established methods exist for assessing hip joint laxity or for the quantitative measure of hip joint stability exist for small animal models. Future development of methods for assessing hip joint instability would benefit the field of hip biomechanics and could be used to identify the joint-level health of the developing or injured hip. A second limitation to this study is that, due to the extent of morphological changes observed after severe instability injury, generalizations of shape variability with such a small sample size may not be appropriate. However, the insight provided does indicate morphological changes and motivates future research.

## Acknowledgements

We acknowledge Patrick Canning (University of Delaware) for his assistance with dynamic histomorphometry and imaging and Daniel Leib (Washington University) for his assistance with microCT image processing. This research was supported by the Washington University Musculoskeletal Research Center (NIH P30 AR057235) and the National Institute of General Medical Sciences of the National Institutes of Health (P41-GM103545, R01-GM083925), and the National Institute of Biomedical Imaging and Bioengineering (R01-EB016701 to AEA). The content is solely the responsibility of the authors and does not necessarily represent the official views of the National Institutes of Health.

## References

1 Lievense, A., Bierma-Zeinstra, S., Verhagen, A., Verhaar, J. & Koes, B. Influence of hip dysplasia on the development of osteoarthritis of the hip. Annals of Rheumatic Diseases 63, 621–626 (2004).

2 Mahan, S. & Kasser, J. Does swaddling influence developmental dysplasia of the hip? Pediatrics 121, 177–178 (2008).

3 Noordin, S., Umer, M., Hafeez, K. & Nawaz, H. Developmental dysplasia of the hip. Orthop Rev (Pavia) 2, e19 (2010).

4 Tanzer, M. & Noiseux, N. Osseous abnormalities and early osteoarthritis: the role of hip impingement. Clinical Orthopaedics and Related Research^®^ 429, 170–177 (2004).

5 Leunig, M., Podeszwa, D., Beck, M., Werlen, S. & Ganz, R. Magnetic resonance arthrography of labral disorders in hips with dysplasia and impingement. Clinical orthopaedics and related research 418, 74–80 (2004).

6 Harris-Hayes, M. & Royer, N. K. Relationship of acetabular dysplasia and femoroacetabular impingement to hip osteoarthritis: a focused review. PM&R 3, 1055–1067 (2011).

7 Barros, H. J. M., Camanho, G. L., Bernabé, A. C., Rodrigues, M. B. & Leme, L. E. G. Femoral head-neck junction deformity is related to osteoarthritis of the hip. Clinical Orthopaedics and Related Research^®^ 468, 1920–1925 (2010).

8 Gala, L., Clohisy, J. & Beaule, P. Hip dysplasia in the young adult. Journal of Bone & Joint Surgery 98, 63–73 (2016).

9 Wenger, D. R. & Bomar, J. D. Human hip dysplasia: evolution of current treatment concepts. Journal of orthopaedic science 8, 264–271 (2003).

10 Buckley, S. L., Sponseller, P. D. & Magid, D. The acetabulum in congenital and neuromuscular hip instability. Journal of pediatric orthopedics 11, 498–501 (1991).

11 Harris, M. D., Davis, R. S., MacWilliams, B. A., Peters, C. L. & Anderson, A. E. Musculoskeletal Modeling of Acetabular Dysplasia-Kinematics, Muscle and Joint Reaction Forces. 795–796, doi:10.1115/SBC2010-19658 (2010).

12 Abraham, C. L., Knight, S. J., Peters, C. L., Weiss, J. A. & Anderson, A. E. Patient-specific chondrolabral contact mechanics in patients with acetabular dysplasia following treatment with peri-acetabular osteotomy. Osteoarthritis and cartilage 25, 676–684, doi:10.1016/j.joca.2016.11.016.

13 Henak, C. R. et al. Patient-specific analysis of cartilage and labrum mechanics in human hips with acetabular dysplasia. Osteoarthritis and cartilage 22, 210–217, doi:10.1016/j.joca.2013.11.003 (2014).

14 Henak, C. R. et al. Role of the acetabular labrum in load support across the hip joint. Journal of biomechanics 44, 2201–2206 (2011).

15 Ng, K. C. G., Rouhi, G., Lamontagne, M. & Beaulé, P. E. Finite element analysis examining the effects of cam FAI on hip joint mechanical loading using subject-specific geometries during standing and maximum squat. HSS Journal^®^ 8, 206–212 (2012).

16 Jorge, J. P. et al. Finite element simulations of a hip joint with femoroacetabular impingement. Computer methods in biomechanics and biomedical engineering 17, 1275–1284 (2014).

17 Ng, K. C. G., Lamontagne, M., Labrosse, M. R. & Beaulé, P. E. Hip joint stresses due to cam-type femoroacetabular impingement: A systematic review of finite element simulations. PloS one 11, e0147813 (2016).

18 Lee, M. & Eberson, C. Growth and development of the child’s hip. Orthopedic Clinics of North America 37, 119–132 (2006).

19 Steppacher, S., Tannast, M., Werlen, S. & Siebenrock, K. Femoral morphology differs between deficient and excessive acetabular coverage. Clinical Orthopaedics and Related Research 466, 782–790 (2008).

20 Clohisy, J., Nunley, R., Carlisle, J. & Schoenecker, P. Incidence and characteristics of femoral deformities in the dysplastic hip.. Clinical Orthopaedics and Related Research 467, 128–134 (2009).

21 Siebenrock, K. A. et al. Abnormal extension of the femoral head epiphysis as a cause of cam impingement. Clinical Orthopaedics and Related Research^®^ 418, 54–60 (2004).

22 Beck, M., Kalhor, M., Leunig, M. & Ganz, R. Hip morphology influences the pattern of damage to the acetabular cartilage: femoroacetabular impingement as a cause of early osteoarthritis of the hip. Bone & Joint Journal 87, 1012–1018 (2005).

23 Hansen, B. J. et al. Correlation between radiographic measures of acetabular morphology with 3D femoral head coverage in patients with acetabular retroversion. Acta Orthopaedica 83, 233–239, doi:10.3109/17453674.2012.684138 (2012).

24 Giorgi, M., Carriero, A., Shefelbine, S. & Nowlan, N. Mechanobiological simulations of prenatal joint morphogenesis. Journal of Biomechanics 47, 989–995 (2014).

25 Nowlan, N., Chandaria, V. & Sharpe, J. Immobilized chicks as a model system for early-onset developmental dysplasia of the hip. Journal of Orthopedic Research 23, 777–785 (2014).

26 Brenner, E., Gruber, H. & Fritsch, H. Fetal development of the first metatarsophalangeal joint complex with special reference to the intersesamoidal ridge. Ann Anat 184, 481–487 (2002).

27 Storm, E. E. & Kingsley, D. M. Joint patterning defects caused by single and double mutations in members of the bone morphogenetic protein (BMP) family. Development 122, 3969–3979 (1996).

28 Ford, C., Nowlan, N., Thomopoulos, S. & Killian, M. The effects of imbalanced muscle loading on hip joint development and maturation. Journal of Orthopaedic Research 35, 1128–1136 (2017).

29 Shefelbine, S. J. & Carter, D. R. Mechanobiological predictions of growth front morphology in developmental hip dysplasia. Journal of Orthopaedic Research 22, 346–352, doi:10.1016/j.orthres.2003.08.004 (2004).

30 Ráliš, Z. & McKibbin, B. Changes in shape of the human hip joint during its development and their relation to its stability. Journal of Bone & Joint Surgery, British Volume 55, 780–785 (1973).

31 Serrat, M., Reno, P., McCollum, M., Meindl, R. & Lovejoy, C. Variation in mammalian proximal femoral development: comparative analysis of two distinct ossification patterns. Journal of Anatomy 210, 249–258 (2007).

32 Hingsammer, A., Bixby, S., Zurakowski, D., Yen, Y.-M. & Kim, Y.-J. How do acetabular version and femoral head coverage change with skeletal maturity? Clinical Orthopaedics and Related Research 473, 1224–1233 (2015).

33 Schindelin, J. et al. Fiji: an open-source platform for biological-image analysis. Nature Methods 9, 676–682 (2012).

34 Doube, M. et al. BoneJ: Free and extensible bone image analysis in ImageJ. Bone 47, 1076–1079 (2010).

35 Cates, J., Fletcher, P., Styner, M., Shenton, M. & Whitaker, R. Shape modeling and analysis with entropy-based particle systems.. Inf Process Med Imaging 20, 333–345 (2007).

36 Maas, S. A., Ellis, B. J., Ateshian, G. A. & Weiss, J. A. FEBio: finite elements for biomechanics. Journal of biomechanical engineering 134, 011005, doi:10.1115/1.4005694 (2012).

37 Team, R. C. R: A language and environment for statistical computing., <http://www.r-project.org/> (2014).

38 Ma, H. L. et al. Osteoarthritis severity is sex dependent in a surgical mouse model. Osteoarthritis and cartilage 15, 695–700, doi:10.1016/j.joca.2006.11.005 (2007).

39 Hashimoto, S., Rai, M. F., Janiszak, K. L., Cheverud, J. M. & Sandell, L. J. Cartilage and bone changes during development of post-traumatic osteoarthritis in selected LGXSM recombinant inbred mice. Osteoarthritis Cartilage 20, 562–571, doi:10.1016/j.joca.2012.01.022 (2012).

40 Christiansen, B. A. et al. Musculoskeletal changes following non-invasive knee injury using a novel mouse model of post-traumatic osteoarthritis. Osteoarthritis Cartilage 20, 773–782, doi:10.1016/j.joca.2012.04.014 (2012).

41 Wang, E. et al. Does swaddling influence developmental dysplasia of the hip?: An experimental study of the traditional straight-leg swaddling model in neonatal rats. The Journal of bone and joint surgery. American volume 94, 1071–1077, doi:10.2106/jbjs.k.00720 (2012).

42 Vanden Berg-Foels, W. S., Todhunter, R. J., Schwager, S. J. & Reeves, A. P. Effect of early postnatal body weight on femoral head ossification onset and hip osteoarthritis in a canine model of developmental dysplasia of the hip. Pediatric research 60, 549–554, doi:10.1203/01.pdr.0000243546.97830.a0 (2006).

43 Wu, W. et al. Sites of collagenase cleavage and denaturation of type II collagen in aging and osteoarthritic articular cartilage and their relationship to the distribution of matrix metalloproteinase 1 and matrix metalloproteinase 13. Arthritis & Rheumatology 46, 2087–2094 (2002).

44 Rousseau, J.-C. & Delmas, P. D. Biological markers in osteoarthritis. Nature Reviews Rheumatology 3, 346 (2007).

45 Henak, C. R. et al. Finite element predictions of cartilage contact mechanics in hips with retroverted acetabula. Osteoarthritis and cartilage 21, 1522–1529 (2013).

46 Harris, M. D. et al. Statistical shape modeling of cam femoroacetabular impingement. Journal of Orthopaedic Research 31, 1620–1626 (2013).

47 Chan, E. F., Farnsworth, C. L., Koziol, J. A., Hosalkar, H. S. & Sah, R. L. Statistical shape modeling of proximal femoral shape deformities in Legg–Calvé–Perthes disease and slipped capital femoral epiphysis. Osteoarthritis and cartilage 21, 443–449, doi:https://doi.org/10.1016/j.joca.2012.12.007 (2013).

48 Chan, E. F. et al. Structural and Functional Maturation of Distal Femoral Cartilage and Bone during Postnatal Development and Growth in Humans and Mice. The Orthopedic Clinics of North America 43, 173–185, doi:10.1016/j.ocl.2012.01.005 (2012).

49 Wang, V. M., Banack, T. M., Tsai, C. W., Flatow, E. L. & Jepsen, K. J. Variability in tendon and knee joint biomechanics among inbred mouse strains. Journal of Orthopaedic Research 24, 1200–1207, doi:10.1002/jor.20167 (2006).

50 Blankevoort, L., van Osch, G. J. V. M., Janssen, B. & Hekman, E. E. G. In vitro laxity-testers for knee joints of mice. Journal of Biomechanics 29, 799–806, doi:http://dx.doi.org/10.1016/0021-9290(95)00130-1 (1996).

